# Host cell glycosylation selects for infection with CCR5- versus CXCR4-tropic HIV-1

**DOI:** 10.1101/2023.09.05.556399

**Authors:** Hannah L. Itell, Daryl Humes, Nell E. Baumgarten, Julie Overbaugh

## Abstract

HIV-1 infection involves a selection bottleneck that leads to transmission of one or a few HIV variants, which nearly always use CCR5 as the coreceptor (R5 viruses) for viral entry as opposed to CXCR4 (X4 viruses). The host properties that drive this selection are not well understood and may hold keys to factors that govern HIV susceptibility. In this report, we identified SLC35A2, a transporter of UDP-galactose, as a candidate X4-specific restriction factor in CRISPR-knockout screens in primary target CD4^+^ T cells. SLC35A2 inactivation in CD4^+^ T cells, which resulted in truncation of glycans due to the absence of galactose, not only increased X4 infection levels, but also consistently decreased infection levels of R5 HIV strains. Single cycle infections demonstrated that the effect is host cell dependent. SLC35A2 is expressed in CD4^+^ T cells at different tissue sites, with high levels in the genital tract – the site of most HIV infections. These data support a role for a host cell protein that regulates glycan structure on HIV infection, with enhanced R5 infection but reduced X4 infection associated with SLC35A2-mediated glycosylation. Host cell glycosylation may therefore contribute to R5 selection and host susceptibility during HIV transmission.

## MAIN TEXT

HIV-1 undergoes a severe viral population bottleneck during transmission. HIV’s high mutation rate coupled with host selective pressures leads to a large diversity of viral variants circulating during chronic infection, yet only one or a few viral variants are detected soon after infection^1-6^. While transmitted viruses are thought to be successful in part due to their high replicative fitness^7,8^ and reduced susceptibility to interferon (IFN)-mediated inhibition^9,10^, the most consistent, defining feature of transmitted variants is their strong coreceptor tropism preferences. HIV can use either CCR5 or CXCR4 as a coreceptor for viral entry^11^, yet transmitted variants nearly always use CCR5^6,12^ despite broad expression of CXCR4 at transmission sites^13^, with CXCR4-tropic (X4) viruses arising in the months to years after primary infection^14^. It has remained a longstanding question in the HIV field why CCR5-tropic (R5) variants, but not X4 viruses, can successfully navigate this bottleneck, leading to new infections. In this study, we aimed to identify host factors that preferentially inhibit X4 HIV, as these may represent selective pressures that contribute to R5 selection during HIV transmission and thus to HIV susceptibility.

To identify X4-specific restriction factors, we performed CRISPR-knockout (KO) screens in the main target cells of HIV infection *in vivo*, primary CD4^+^ T cells, and infected with an X4 strain (LAI^15^) and an R5 virus (Q23.BG505^16^) for comparison. We leveraged the HIV-CRISPR screening strategy^17^, which we recently adapted for use in primary T cells^18^, and interrogated ∼2,000 host genes, including many IFN-stimulated genes (ISGs) as they are a major driver of antiviral immunity (**Fig. 1A; Supplementary Table 1**). Screens were therefore performed in the context of type I IFN treatment. Genes required for IFN signaling were enriched for all screens and we observed strong agreement between screens using primary T cells from different donors (**Fig. 1B; Supplementary Table 2**). By comparing Q23.BG505 and LAI screens and selecting LAI-specific hits, we identified 81 candidate X4-specific restriction factors, of which the top-scoring hit was SLC35A2 (**Fig. 1C**). This gene was of particular interest because it was highly enriched for both LAI screens and conversely scored below background for both Q23.BG505 screens (**Fig. 1D**). SLC35A2 was not upregulated by IFN in primary CD4^+^ T cells (**Fig. 1E**). These data support SLC35A2’s candidacy as a non-ISG host factor that may preferentially inhibit X4 HIV.

**Figure 1.**
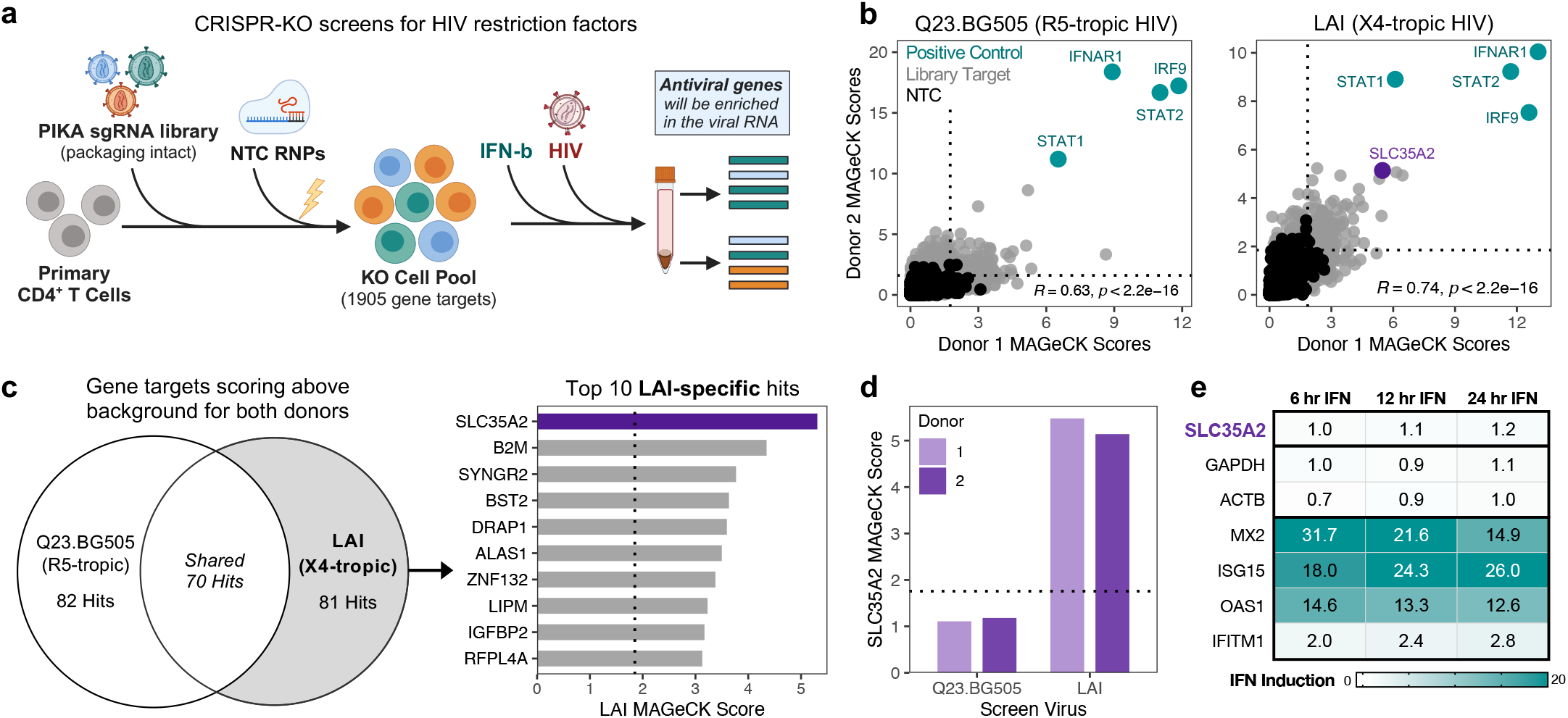
SLC35A2 is an X4-specific hit in a CRISPR-KO screen for HIV restriction factors in primary CD4^+^ T cells. **A**. Schematic of HIV-CRISPR screening approach. **B**. Correlation of positive MAGeCK scores between CD4^+^ T cell donors in Q23.BG505 (left) and LAI (right) screens. Pearson correlation results are included within each plot. Dotted lines denote background for each screen based on non-targeting control (NTC) results (background = average NTC + 3 standard deviations). **C**. Left, Euler diagram comparing hits between Q23.BG505 and LAI screens. Screen hits are defined as gene targets that scored above background for both donors (top right quadrants from Panel B). Right, top 10 scoring LAI-specific hits, ordered by their average MAGeCK score in LAI screens. Dotted line reflects average background from LAI screens. **D**. MAGeCK scores for SLC35A2 in each HIV-CRISPR screen. Dotted line indicates average background for all screens. **E**. Heatmap of average IFN-induced fold changes in RNA expression for SLC35A2, housekeeping genes (GAPDH, ACTB), and canonical ISGs (MX2, ISG15, OAS1, IFITM1) in primary CD4^+^ T cells at 6-, 12-, and 24-hours post-IFN treatment (n = 3, 3, and 5 donors, respectively by time point).

To validate our CRISPR-KO screen findings, we performed single gene knockouts in primary CD4^+^ T cells, inactivating SLC35A2 and the B cell marker CD19 as a negative control. Across several unique donors and independent experiments, we found that SLC35A2 KO not only significantly increased infection levels of X4-tropic LAI (5.9-fold decrease, p=0.004), but, surprisingly, it also significantly decreased those of R5-tropic Q23.BG505 (4.8-fold increase, p<0.0001) (**Fig. 2A**). These findings were consistent for two separate assays of HIV infection: quantifying reverse transcriptase (RT) activity in infection supernatants (**Fig. 2A, Extended Data Fig. 1A**) and measuring the percentage of cells staining positive for HIV-Gag (**Fig. 2B, Extended Data Fig. 1B, C**). Notably, based on Gag staining, an average of 73% of SLC35A2 KO cells across four unique donors (range: 64.8-83.2%) were infected with LAI compared to 13% in CD19 KO control cells (range: 9.6-16%) (5.9-fold decrease, p=0.001) (**Fig. 2B, Extended Data Fig. 1C**). The opposite trend was true for Q23.BG505, with 1.8% of SLC35A2 KO cells (range: 1-2.8%) staining positive for HIV-Gag compared to 11% of CD19 KO cells (range: 7.8-16.4%) (7.6-fold increase, p=0.03). SLC35A2 KO therefore causes substantial, opposing phenotypes for these two HIV strains that differ in their coreceptor usage.

**Figure 2.**
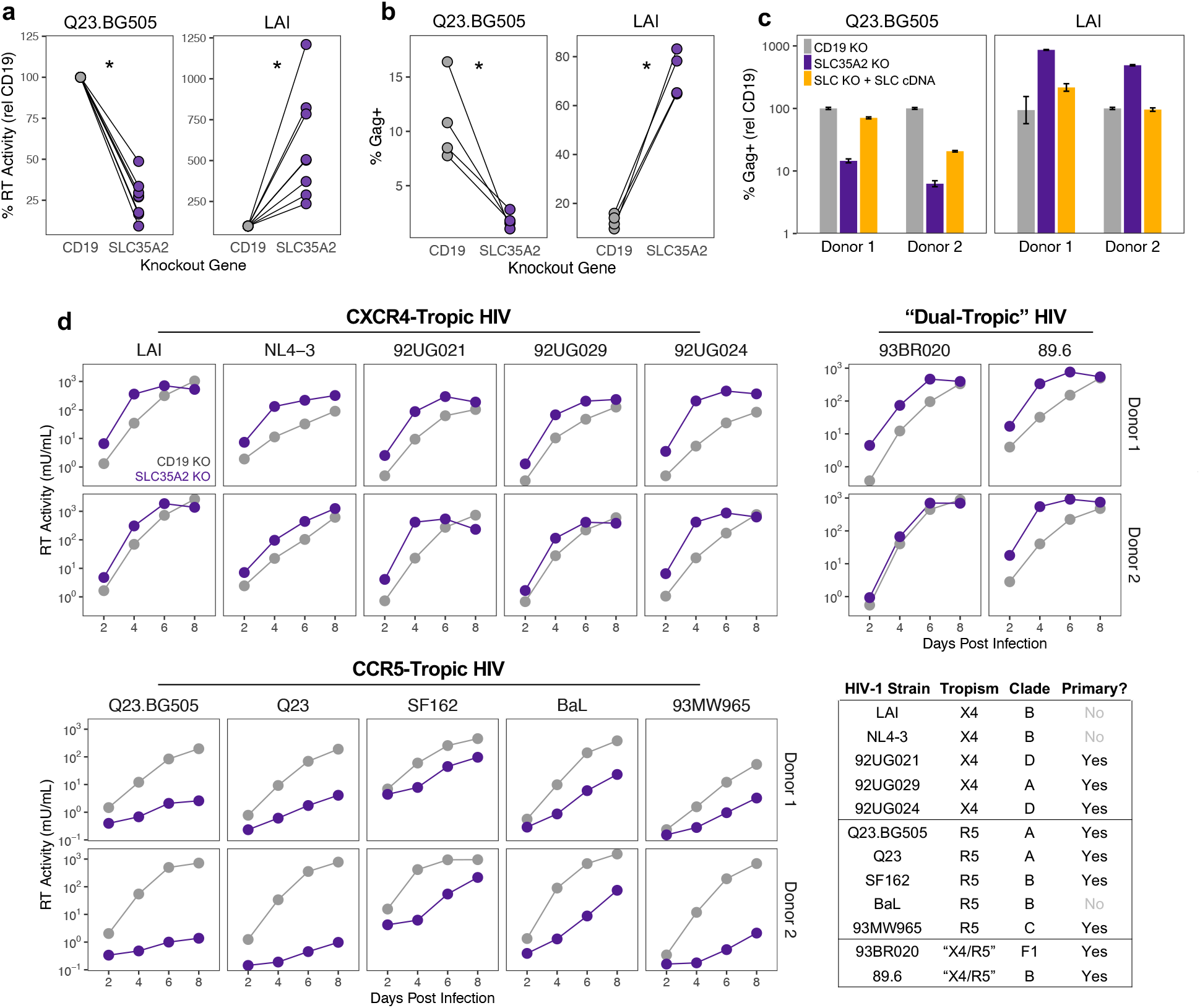
SLC35A2 KO differentially impacts CXCR4-tropic and CCR5-tropic HIV. **A**. Reverse transcriptase (RT) activity relative to CD19 KO at 2 days post infection (dpi) (MOI=0.02). Knockouts were performed in CD4^+^ T cells from five donors across single or replicate knockout and infection experiments for a total of 8 independent experiments. Editing efficiency ranged from 52% to 96% KO (82% KO average, n=8). Two-tailed paired t-test, asterisk indicates p<0.05. **B**. Percentage of CD4^+^ T cells staining positive for HIV-Gag at 3 dpi (MOI=1) (n=4 independent donors). Two-tailed paired t-test, asterisk indicates p<0.05. **C**. Percentage of CD4^+^ T cells staining positive for HIV-Gag at 3 dpi (MOI=1), relative to CD19 KO. Data are shown as mean ± standard deviation of duplicate infections. **D**. RT activity over time from spreading infections (MOI=0.02) in primary CD4^+^ T cells from two donors. Results are separated based on coreceptor tropism. Bottom right table, characteristics of the HIV strains used for infections.

To determine whether the observed phenotypes were specific to SLC35A2 inactivation and not due to off-target effects, we exogenously expressed SLC35A2 cDNA in SLC35A2 KO CD4^+^ T cells. Complementation led to full or partial rescue of wildtype HIV infection levels (**Fig. 2C**). Cases of partial rescue are likely due to the polyclonal nature of both SLC35A2 KO and complementation, as these primary cell culture experiments are not amenable to antibiotic or clonal selection. These data support the specificity of SLC35A2 KO.

Though the opposing SLC35A2 KO phenotypes for Q23.BG505 and LAI were consistent across experimental platforms, donors, and infection metrics, it is unclear what biological viral characteristics are driving these effects. One major difference between these viruses is their coreceptor usage; however, they are also from different HIV clades and LAI is lab-adapted whereas Q23.BG505 is not. To understand which viral features represent determinants of the phenotype of SLC35A2 KO on HIV infection, we infected SLC35A2 KO and CD19 KO CD4^+^ T cells with five X4 and five R5 full-length HIV strains from various clades and consisting of both primary and lab-adapted viruses. We observed a strikingly consistent pattern – SLC35A2 KO increased infection levels for all X4 strains whereas all R5 viruses exhibited lower levels of infection (**Fig. 2D, Extended Data Fig. 2A**). Coreceptor tropism for each strain was confirmed by individually inactivating CXCR4 and CCR5 in CD4^+^ T cells (**Extended Data Fig. 2B**). In addition to X4 and R5 HIV, we investigated the effect of SLC35A2 KO on two viruses considered dual-tropic (93BR020^19^ and 89.6^20^). These results closely recapitulated those of X4 HIV (**Fig. 2D, Extended Data Fig. 2A**), which agrees with our findings from coreceptor KO cells that indicated that both 93BR020 and 89.6 rely on CXCR4 substantially more than CCR5 (**Extended Data Fig. 2B**). Inactivation of SLC35A2 therefore differentially impacts HIV based on whether viruses use CXCR4 or CCR5 as a coreceptor for viral entry.

SLC35A2 is a multichannel membrane protein that transports UDP-galactose from the cytosol into the Golgi^21,22^. This process is critical for normal glycosylation as galactose is a common residue on both N- and O-glycans. Therefore, SLC35A2 inactivation results in truncated glycans that lack galactose^23^. To determine whether SLC35A2 KO disrupts glycosylation in our primary cell system, we stained SLC35A2 KO CD4^+^ T cells with two lectins that bind terminal glycan residues that would be expected to be exposed with the loss of galactose (**Fig. 3A**). Specifically, *Griffonia Simplicifolia* lectin II (GSL-II) preferentially binds terminal GlcNAcs on N-glycans and *Vicia Villosa* lectin (VVL) recognizes terminal GalNAcs, which are residues are specific to O-glycans^24^. We found that 94% and 91% of SLC35A2 KO cells were positive for GSL-II and VVL binding, respectively, compared to 3% and 1% of CD19 KO cells (**Fig. 3B**). Therefore, despite the polyclonal nature of our KOs, editing of SLC35A2 in this cell type substantially disrupts host cell surface glycosylation by causing the truncation of both N- and O-glycans.

**Figure 3.**
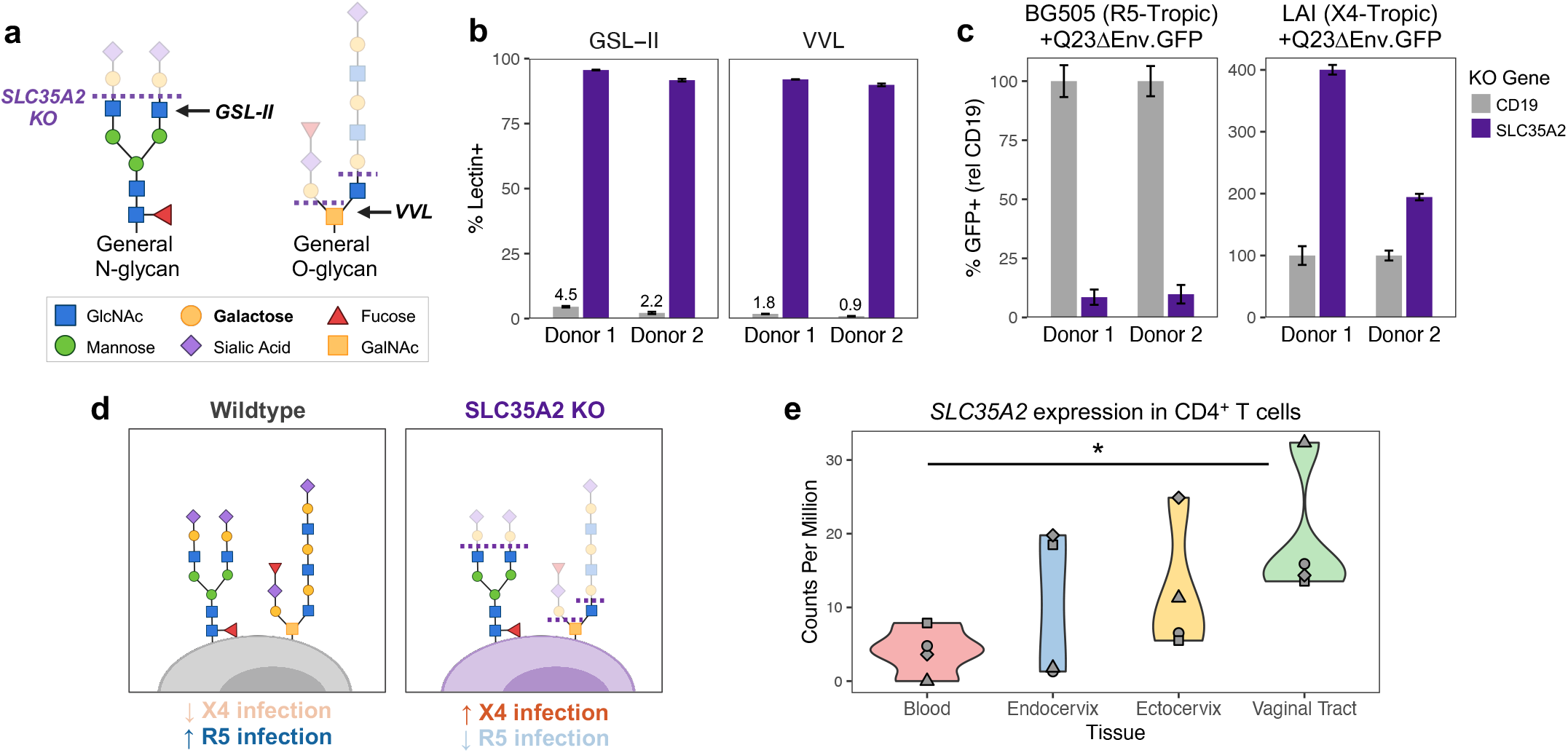
SLC35A2 inactivation causes truncated glycans on host cells and impacts HIV infection within a single round of infection. **A**. General schematic of glycans, including the expected truncation when SLC35A2 is inactivated. The terminal residue binding sites of GSL-II and VVL lectins are indicated with arrows. **B**. Percentage of cells staining positive for GSL-II and VVL lectins in CD4^+^ T cells from two donors. Data are shown as mean ± standard deviation of 2-3 replicate stains. **C**. Percentage of GFP+ CD4^+^ T cells, relative to CD19 KO, after 2 days of infection with GFP-expressing HIV pseudoviruses (MOI=1). Data are shown as mean ± standard deviation of 3 replicate infections. **D**. Working model of the impact of CD4^+^ T cell glycosylation on HIV infection. **E**. *SLC35A2* expression in CD4^+^ T cells isolated from different anatomical compartments from four donors. Shapes denote donors. One-tailed paired t-test, asterisk indicates p<0.05.

Though lectin staining demonstrates altered glycosylation on target cells, HIV itself is heavily glycosylated and viral glycans are likely similarly altered upon replication in SLC35A2 KO cells. Because our infections have thus far utilized replication-competent HIV, the observed tropism-specific SLC35A2 KO HIV phenotypes may be driven by altered glycans either on host cells or on HIV virions after at least one round of replication in SLC35A2 KO cells. To distinguish these possibilities, we made X4 and R5 HIV pseudoviruses, which are capable of infection but not making infectious progeny virions. We found that SLC35A2 KO impacted these HIV pseudoviruses within a single infection cycle in the same manner as replication-competent virus, with a reduction in R5 infection and an increase in X4 virus infection (**Fig. 3C**). The magnitudes of these effects (BG505 average: 10.9-fold decrease, LAI average: 3-fold increase) were similar to those from replication-competent infection (**Fig. 2B**). The effect of SLC35A2 KO on HIV is therefore driven by changes to glycans on target cells, not on progeny virus.

These data support an opposing, early role for CD4^+^ T cell glycans on R5 and X4 HIV infection, whereby R5 HIV prefers wildtype target cell glycosylation and X4 viruses are more successful when host glycans are truncated, such as in the setting of SLC35A2 KO (**Fig. 3D**). Interestingly, these results may explain the cellular driver for recent observations by Ma *et al*. showing that memory CD4^+^ T cells from the blood and tonsils that have higher levels of binding to WGA, a lectin that binds sialic acid and GlcNAc, are more susceptible to R5 HIV infection, whereas removal of sialic acid reduced R5 infection^25^. Together, our findings suggest that sialic acid residues, which are absent in the setting of SLC35A2 KO, may specifically potentiate R5 HIV infection of CD4^+^ T cells and identify a specific host gene central to this process.

Our results not only provide insight into determinants of R5 infection; they also uniquely demonstrate that target cell glycan features differentially impact HIV infection based on coreceptor tropism. Based on the existing literature and our working model (**Fig. 3D**), if host glycans were to influence R5 selection during HIV transmission, we would expect there to be high levels of SLC35A2 and sialic acid on target cells that HIV first encounters in the mucosa. To address this, we analyzed a previously published bulk RNA-seq dataset of CD4^+^ T cells isolated from the blood or three female genital tract sites^26^ and found that *SLC35A2* was expressed in all CD4^+^ T cell populations, with elevated levels in CD4^+^ T cells from the vaginal tract, a common site of HIV transmission, as compared those in the blood (**Fig. 3E**). Notably, CXCR4 was highly expressed in CD4^+^ T cells from all tissue sites in this dataset, particularly in mucosal tissues, consistent with previous findings^13^ that an absence of CXCR4 expression on relevant target cells does not explain R5 selection during transmission (**Extended Data Figure 3**). In accordance with the *SLC35A2* expression level trends, memory CD4^+^ T cells from the endometrium were reported to have higher levels of WGA lectin binding (sialic acid/GlcNAc) than those from the blood^25^. Target cells in HIV transmission sites therefore have SLC35A2 levels and glycan features that support infection of R5 viruses and may restrict that of X4 viruses. Host glycosylation may therefore contribute to R5 selection during the HIV transmission bottleneck.

It remains to be determined whether the opposing tropism phenotypes with SLC35A2 KO are due to changes to specific glycans on target cells, such as glycans on the coreceptors themselves, or are due to broad changes in the overall glycan landscape of target cells. CXCR4 and CCR5 both harbor glycans on their extracellular N-termini^27-29^, and this region of CCR5 is known to interact with HIV^30^. It is therefore possible that the impact of SLC35A2 is on the composition and structure of glycans on the coreceptors, with subsequent coreceptor-dependent effects on HIV binding and/or fusion. However, CD4^+^ T cells express a variety of glycoproteins and a recent study concluded that general glycan-glycan interactions between HIV and the host cell may enhance attachment^31^. These glycan-glycan interactions could also be a mechanism through which SLC35A2 impacts HIV infection, though there is at present no evidence for a tropism-specific effect with these broad glycan-glycan interactions. Overall, our findings provide a window into the mechanism behind the tropism-dependent impact of target cell glycosylation on HIV. Further, they address a long-standing question in the field that may inform the development of new prophylactic approaches to specifically target viral variants that are successful in the HIV transmission bottleneck and thus a key to HIV susceptibility.

## EXTENDED DATA FIGURES

**Extended Data Figure 1.**
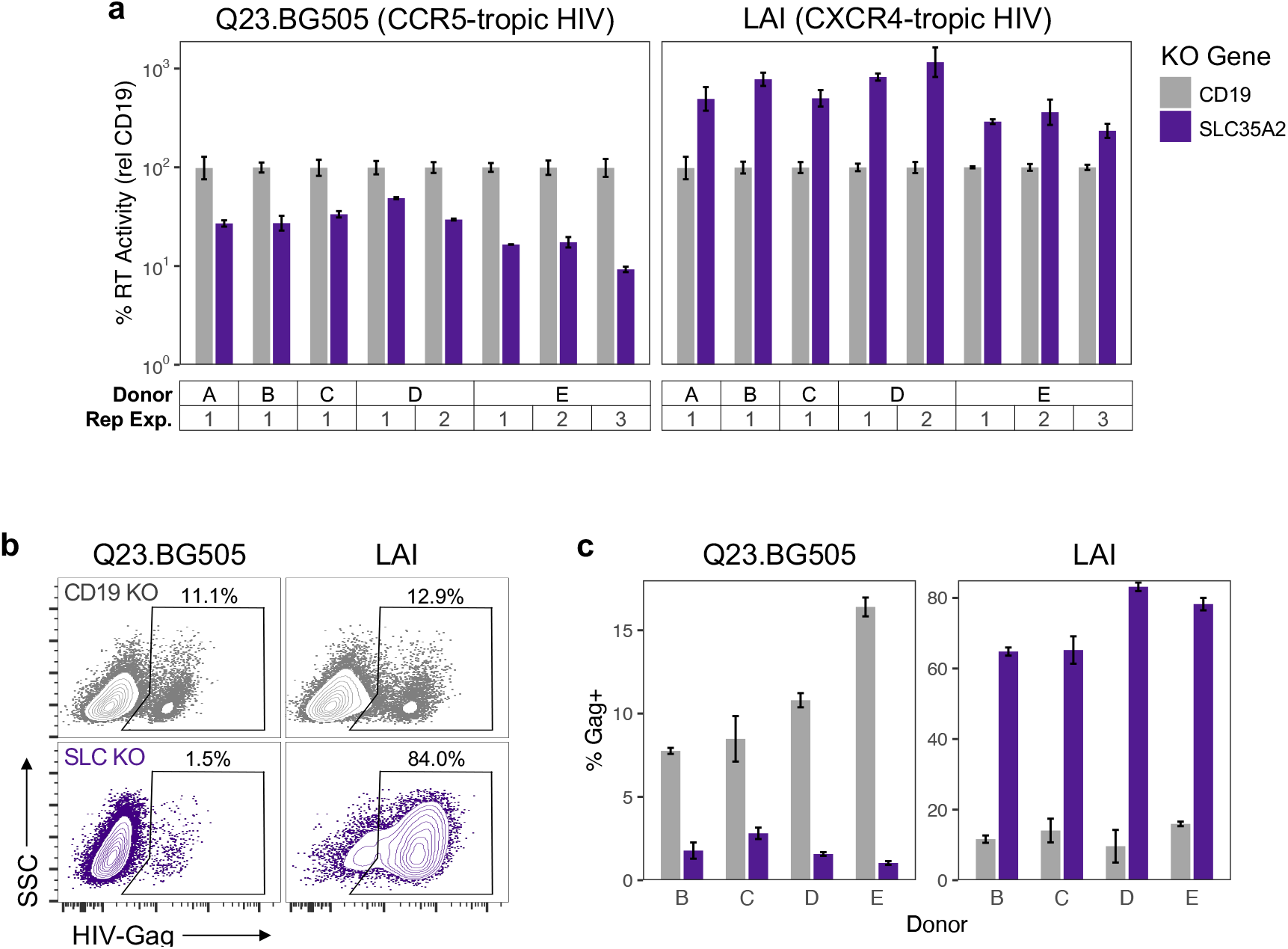
SLC35A2 KO has opposite effects on two HIV strains that utilize different coreceptors, related to Figure 2A and 2B. **A**. Technical and donor replicates for data depicted in Figure 2A. Reverse transcriptase (RT) activity, relative to CD19 KO for each primary CD4^+^ T cell donor/experiment, at 2 days post infection (dpi) (MOI=0.02). Knockouts were performed in five donors across independent replicate KO and infection experiments where indicated. Data are shown as mean ± standard deviation of 2-3 replicate infections. **B**. Representative flow plots of HIV-Gag-FITC staining CD4^+^ T cells from one donor after 3 dpi (MOI=1). **C**. Technical and donor replicates for data depicted in Figure 2B. Percentage of CD4^+^ T cells staining positive for HIV-Gag-FITC at 3 dpi (MOI=1). Donor letters correspond with those in Panel A. Data are shown as mean ± standard deviation of 2-3 replicate infections.

**Extended Data Figure 2.**
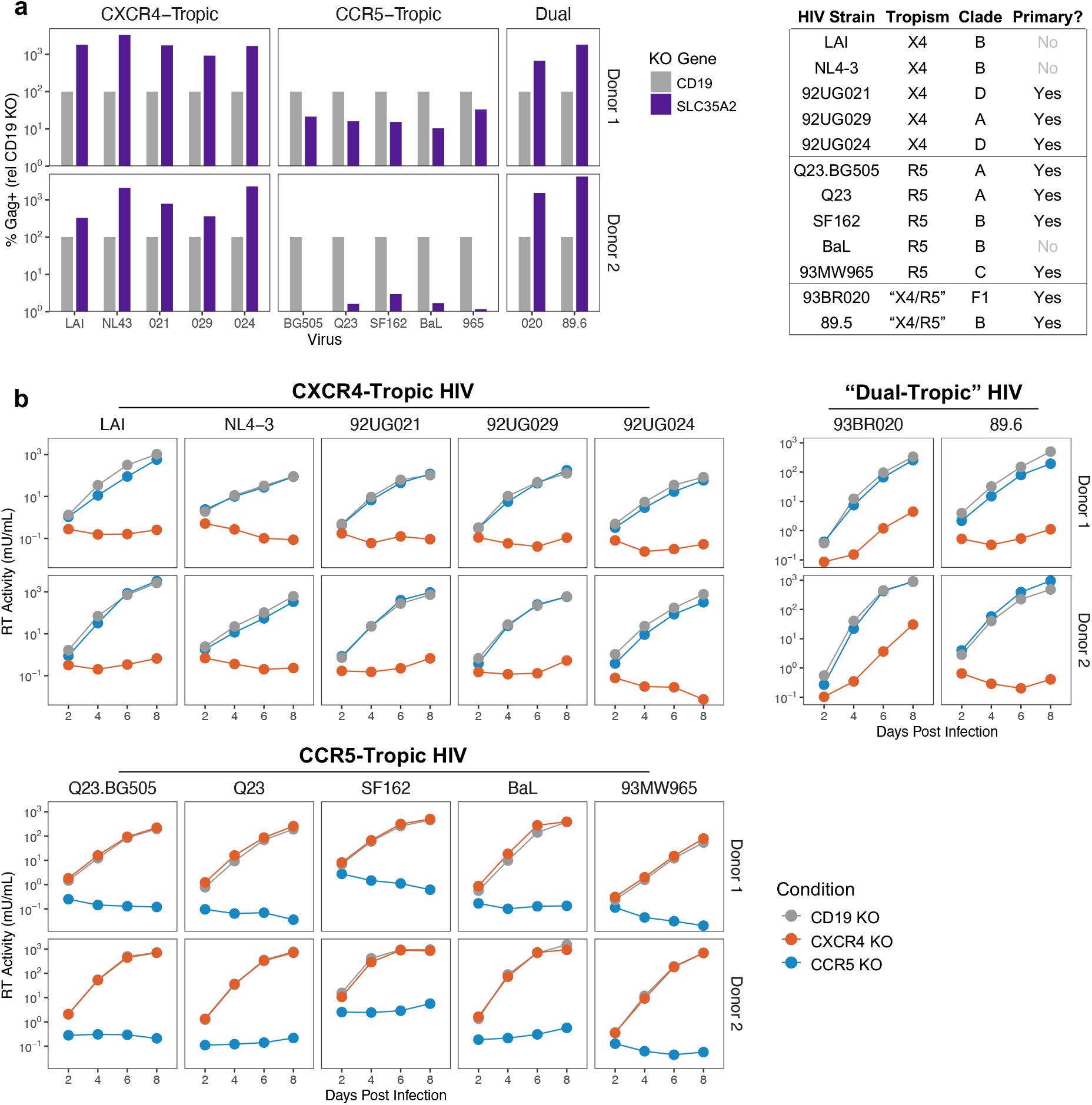
SLC35A2 KO differentially impacts CXCR4-tropic and CCR5-tropic HIV, related to Figure 2D. **A**. Percentage of CD4^+^ T cells staining positive for HIV-Gag, relative to CD19 KO, at 6 dpi (MOI=0.02, same infections as depicted in Figure 2A, different measurement of HIV infection). Donor numbering is consistent with Figure 2D. **B**. RT activity over time from spreading infections (MOI=0.02) in primary CD4^+^ T cells from two donors to determine coreceptor tropism. Results are separated by expected coreceptor tropism based on prior literature. Donor numbering is consistent with Panel A and Figure 2D.

**Extended Data Figure 3.**
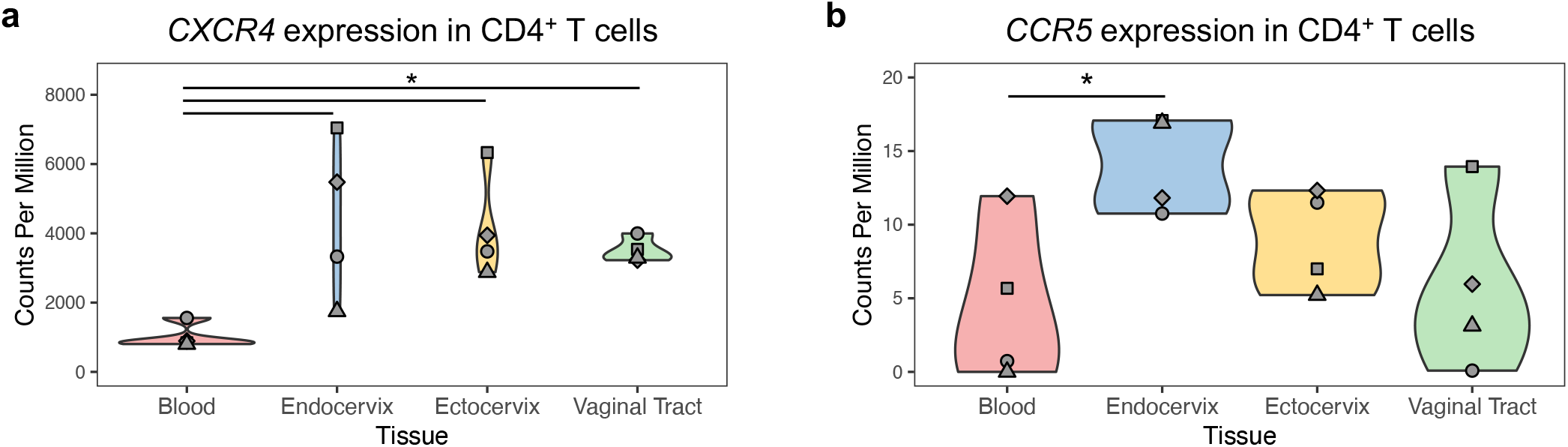
Expression of HIV coreceptors in CD4^+^ T cells from the blood and common HIV transmission sites. (**A**) *CXCR4* and (**B**) *CCR5* expression in CD4^+^ T cells isolated from different anatomical compartments from four donors. Shapes denote donors. One-tailed paired t-test, asterisk indicates p<0.05.

## SUPPLEMENTARY INFORMATION

**Supplementary Table 1**. PIKA guide library with synthetic NTC gene assignments.

**Supplementary Table 2**. HIV-CRISPR screen results.

**Supplementary Table 3**. Guide sequences for single-gene editing experiments.

## METHODS

### Cell lines

HEK293T/17 (ATCC #CRL-11268) and TZM-bl cells (NIH HIV Reagent Program #ARP-8129) were cultured in DMEM complete media (DMEM (Gibco #11965092) supplemented with 10% FBS (Gibco #26140079), 2 mM L-glutamine (Gibco #25030081), 1X antibiotic-antimycotic (Gibco #15240062) that contains penicillin, streptomycin, and Amphotericin B).

### HIV-1 viruses

HIV Q23.BG505^16^ is a clade A, CCR5-tropic HIV chimeric virus derived from a full-length provirus isolated early in infection (Q23^32^) and bears the BG505.C2 envelope^33^ of a well-studied clade A transmitted/founder virus. HIV LAI is a clade B, CXCR4-tropic HIV strain, as previously described^15^. Infectious molecular clones (IMCs) of Q23.BG505^16^ and LAI^15^, as well as Q23^32^, SF162^34^, and NL4-3^35^ (NIH HIV Reagent Program ARP-2852, contributed by Dr. M. Martin) were generated by transfection using proviral plasmids. The remaining viruses were obtained as viral stocks through the NIH HIV Reagent Program, Division of AIDS, NIAID, NIH: HIV 92/UG/021 (ARP-1648), HIV 92/UG/029 (ARP-1650), HIV 92/UG/024 (ARP-1649), and HIV 93/BR/020 (ARP-2329) contributed by UNAIDS Network for HIV Isolation and Characterization; HIV Ba-L (ARP-510) contributed by Dr. Suzanne Gartner, Dr. Mikulas Popovic, and Dr. Robert Gallo^36^; HIV 93/MW/965 (ARP-2913) contributed by Dr. Paolo Miotti and the UNAIDS Network for HIV Isolation and Characterization; HIV 89.6 (ARP-1966) contributed by Dr. Ronald Collman^37^.

### HIV-1 production

HIV IMCs and pseudoviruses were prepared by transfecting HEK293T/17 cells in a six-well format. Twenty hours prior to transfection, 5e5 cells/well were plated in 2 mL of DMEM complete. For each HIV IMC transfection well, 1 µg of proviral DNA and 200 µL serum-free DMEM were combined, prior to the addition of 18 µL of FuGENE (Promega #E2692). Pseudoviruses were also made by transfections using plasmids expressing the envelope of BG505.C2^33^ or LAI (gift from Dr. Michael Emerman) and Q23ΔEnv.GFP^38^ (NIH HIV Reagent Program ARP-12647). For each HIV pseudovirus transfection well, 667 µg of envelope DNA, 1.334 µg of Q23ΔEnv.GFP, and 200 µL serum-free DMEM were combined, prior to the addition of 36 µL of FuGENE. Transfection complexes were gently mixed, incubated for 15-30 minutes at room temperature, and added to each well in a drop-wise manner. At 20 h post-transfection, media was replaced with 1.5 mL of DMEM complete per well. Supernatants were harvested two days post-transfection and clarified using a 0.2 µM filter (Millipore Sigma #SCGP00525). Transfection supernatants were concentrated using Amicon 100 KDa concentrators (Millipore Sigma #UFC910024) and titered on TZM-bl cells, as previously described^18^.

### Lentiviral plasmids

Lentiviruses were used to deliver the CRISPR guide library to CD4^+^ T cells for HIV-CRISPR screens and SLC35A2 cDNA to SLC35A2 KO CD4^+^ T cells for rescue experiments. To generate the lentiviral vector for HIV-CRISPR screens, the PIKA guide library was amplified from the HIV-CRISPR PIKA library plasmid^17^ and cloned into the HIV-TOP opt mChΔW^12^ plasmid for primary CD4^+^ T cell screens, as previously described^18^. To generate the lentiviral vector for SLC35A2 rescue experiments, SLC35A2 cDNA (393 aa, NCBI Reference Sequence: NP_001035963.1) was codon-optimized and synthesized by Twist Bioscience, amplified, and cloned into the TOP-CAR expression vector^39^. Specifically, the codon-optimized SLC35A2 fragment, WPRE-LTR from the HIV-CRISPR PIKA plasmid^17^, and AgeI- and PmeI-digested TOP-CAR were assembled using HiFi DNA Assembly Master Mix (NEB #E2621L) to generate the TOP-SLC35A2 expression plasmid.

### Lentivirus production

Lentiviruses were transfected as described above for HIV production, using the following transfection reagents per well: 667 ng of transfer vector (HIV-TOP opt mChΔW with the PIKA guide library or TOP-SLC35A2), 500 ng of psPAX2 packing vector (Addgene #12260), 333 ng of the pMD2.G VSV-G expression vector (Addgene #12259), 200 µL serum-free DMEM, and 4.5 µL TransIT-LT1 transfection reagent (Mirus Bio #MIR-2305). Lentiviral transfections were concentrated over a sucrose gradient, as previously described^18^.

### Preparation of ribonucleoproteins (RNPs)

Non-targeting control (NTC) RNPs utilized a single sgRNA from IDT, whereas RNPs for all other single gene KOs were assembled using a mixture of three sgRNAs from Synthego (Synthego Gene KO Kit v2). Guide sequences are available in **Supplementary Table 3**. RNPs were prepared with Cas9-NLS protein (UC Berkeley QB3 MacroLab) as described previously^18^.

### Overview of primary CD4^+^ T cell experiments

In brief, this study reports three types of primary CD4^+^ T cell editing and infection experiments: (1) single gene editing, (2) HIV-CRISPR screens, and (3) SLC35A2 rescue experiments. All primary CD4^+^ T cell experiments utilize the following methods and timeline, with additional methods for specific experiments as indicated: CD4^+^ T cell isolation on Day 1, CD4^+^ T cell activation on Day 1, transduction of CD4^+^ T cells with lentivirus on Day 2 (*CRISPR screens and SLC35A2 rescue experiments only)*, Cas9 delivery and genomic editing of CD4^+^ T cells by nucleofection on Day 4, and HIV infection on Day 14. This experimental timeline and all related methods were recently described in detail by our group^18^. General protocols, as well as methods specific to CRISPR screens and SLC35A2 rescue experiments, are described in the following five sections.

### Primary CD4^+^ T cell isolation and activation

Peripheral blood mononuclear cells (PBMCs) were isolated from whole blood from healthy donors by centrifugation over Ficoll-Paque Plus (Cytiva #17144002). CD4^+^ T cells were isolated using negative selection (STEMCELL Technologies #17952) and were immediately cultured in complete RPMI media (RPMI 1640 (Gibco #22400089) supplemented with 10% FBS, 2 mM L-glutamine, 1X antibiotic-antimycotic) with 100 U/mL recombinant interleukin (IL)-2 (Roche #11147528001) and activated or frozen in freezing media (90% FBS, 10% DMSO). CD4^+^ T cells were used directly after isolation for HIV-CRISPR screens and were thawed from frozen aliquots for all other experiments. After isolation or thawing, cells were resuspended at 2.5e6 cells/mL in 48-well flat-bottom plates with complete RPMI media and 100 U/mL IL-2. Cells were activated with plate-bound anti-CD3 (Tonbo Biosciences #40-0038) at a concentration of 10 µg/mL and anti-CD28 (Tonbo Biosciences #40-0289) supplemented to the media at a concentration of 5 µg/mL.

### RNP nucleofection of CD4^+^ T cells

Three days after activation (Experiment Day 4), nucleofections were performed using 1e6 CD4^+^ T cells per nucleofection as previously descibed^18^. For nucleofections with NTC RNPs, cells were resuspended in 20 µL of P3 Primary Cell Nucleofector solution (Lonza #V4SP-3096) and mixed gently with 4 µL of NTC complexes. For RNPs with Synthego guides, cells were gently resuspended with 25 µL of RNPs. Nucleofections were carried out in a 16-well Nucleocuvette (Lonza #V4SP-3096) using an Amaxa Nucleofector (Lonza) and pulse code EH-115. Cells were allowed to recover for 30 min to 2 h at 37C and were then brought to 2.5e6 cells/mL in RPMI complete media with 100 U/mL IL-2 and at a 1:1 ratio with activation beads (Miltenyi Biotec #130-091-441). Two days later, 200 µL of RPMI complete media with 100 U/mL IL-2 was added to each nucleofection. At four days post nucleofection, half of the culture media was replaced with fresh RPMI complete media with 100 U/mL IL-2 and cells were brought to 1e6 cells/mL. Starting the following day, cells were counted and resuspended to 1e6 cells/mL in RPMI complete media with 100 U/mL IL-2 every other day.

### HIV infection of CD4^+^ T cells

Ten days post-nucleofection (Experiment Day 14), HIV infections were established via spinoculation at 1100xg for 90 min at 30C in the presence of 8 µg/mL polybrene (Millipore Sigma #TR-1003-G) as previously described^18^. After spinoculation, infection inoculum was replaced with fresh RPMI complete with 100 U/mL IL-2. Viral supernatants were harvested at various days post infection as indicated in figure legends and stored at -80C.

### HIV-CRISPR screening

On the day after CD4^+^ T cell isolation and activation (Experiment Day 2), cells were transduced with HIV-TOP opt mChΔW lentivirus containing the PIKA library by spinoculation in the presence of 8 µg/mL protamine sulfate (Millipore Sigma #P3369). Two days later (Experiment Day 4), Cas9 was delivered to cells by nucleofection with NTC RNPs. Nine days after nucleofection (Experiment Day 4), all library cells were treated with 1000 U/mL IFN-β1a (PBL Assay Science #11410). On the following day, infections with HIV IMCs (MOI=1) were established. Cells were split 1:2 three days post infection with fresh RPMI complete media with 100 U/mL IL-2 and 1000 U/mL IFN. Viral supernatants were harvested 3, 4, and 5 dpi and cells were collected 5 dpi. All viral supernatants were filtered to remove cellular debris (Millipore Sigma #SCGP00525) and stored at 4C up to four days post collection. Viral supernatants across collection days were combined and concentrated over a sucrose gradient, resuspended in 100 µL PBS, and stored at 4C. Cells were washed in PBS, pelleted, and stored at -80C.

Guide sequences were amplified from genomic DNA extracted from cell pellets and viral RNA extracted from concentrated viral pellets and were prepared for Illumina sequencing as previously described^18^. Pooled libraries were sequenced on an Illumina HiSeq 2500 (Fred Hutch Shared Resources Genomics Shared Resource). Illumina sequencing reads (SRA accession number: TBD) were analyzed by MAGeCK-Flute^40^ to generate guide counts and gene-level enrichment data (**Supplementary Table 2**; Fred Hutch Center for Data Visualization). NTC guide sequences were iteratively binned to create a number of NTC “genes” (synthetic NTCs, synNTCs) with eight guides per gene to match the number of genes in the PIKA library (1906 synNTCs total; **Supplementary Table 1**). This approach captures variation across NTC guides when analyzing gene-level data more accurately than aggregating all 200 guides into one NTC gene-level data point. Pearson R correlations between MAGeCK scores from different donors were determined using the stat_cor function in the ggpubr package. Screen background was calculated as the average synNTC MAGeCK score + 3 standard deviations.

### SLC35A2 rescue experiments

Primary CD4^+^ T cells were transduced with TOP-SLC35A2 lentivirus on the day after cells were initially thawed and activated. Transductions were performed via spinoculation at 1100xg for 90 min at 30C in the presence of 8 µg/mL protamine sulfate (Millipore Sigma #P3369). Cells were nucleofected with SLC35A2 RNPs two days later, following the normal timeline of all other gene editing and HIV infection experiments. Importantly, SLC35A2 RNPs are unable to target and edit the exogenous codon-optimized SLC35A2 cDNA delivered via transduction with TOP-SLC35A2 lentivirus.

### Reverse transcriptase (RT) activity qPCR assay

HIV infection levels were assessed by measuring RT activity (mU/mL) in viral supernatants using the previously described RT activity qPCR assay^18,41^. The RT units of a concentrated stock of HIV Q23.BG505 virus were determined multiple times using a standard curve of purified HIV RT enzyme (Worthington Biochemical Corp. #LS05003). Aliquots of this titered stock of Q23.BG505 were used as the quantitative standard curve in all assays. The RT activity (mU/mL) of experimental wells was interpolated from the quantitative standard curve using the lm function in the R stats package.

### HIV-Gag staining

After HIV infection, as indicated in figure legends, cells were collected, fixed in 4% paraformaldehyde (Santa Cruz Biotechnology #sc-281692) for 10 min, and either stored at 4C for up to two days or immediately prepared for staining. Cell pellets were permeabilized for intracellular staining via resuspension in 0.5% Triton X-100 in PBS for 10 min and were subsequently resuspended in a 1:500 dilution of anti-p24-FITC KC57 antibody (Beckman Coulter #6604665) in 1% BSA in PBS for 1 h at room temperature in the dark. Washed cells were analyzed by flow cytometry on a BD LSRFortessa X-50 or BD FACSymphony A5.

### Genomic editing analysis

SLC35A2 KO CD4^+^ T cells were collected on the day of HIV infection to assess for genomic editing as previously described in detail^18^. In brief, regions containing SLC35A2 guide cut sites were amplified from genomic DNA (forward primer: 5’ AAGCTGCCCACAAATAGCCT; reverse primer: 5’ AAGCTGCCCACAAATAGCCT) and sequenced (sequencing primer: 5’ AAGCTGCCCACAAATAGCCT) by Sanger sequencing (Fred Hutch Shared Resources Genomics Core). Chromatograms were analyzed by ICE analysis tool v3 (Synthego) to determine the rate of editing predicted to cause SLC35A2 KO.

### Lectin staining

On the day of HIV infection, 2e5 edited CD4^+^ T cells per stain replicate were collected, fixed in 4% paraformaldehyde for 10 min, and resuspended in 20 µg/mL VVL-FITC (Vector Laboratories #FL-1231-2) and 10 µg/mL GSL-II-AF647 (ThermoFisher #L32451) in 2% FBS in PBS for 1 h at room temperature in the dark. Cells were washed, resuspended in 2% FBS in PBS, and were analyzed by flow cytometry on a BD LSRFortessa X-50 or BD FACSymphony A5.

### Analysis of IFN-inducibility

We previously reported a bulk RNA-seq dataset of primary CD4^+^ T cells treated with IFN-β1a (SRA: PRJNA921704)^18^. Here, we analyzed this dataset to determine the fold induction of a subset of genes at various times post IFN treatment. Fold inductions were averaged across unique donors (3 donors for 6 and 12 h timepoints, 5 donors for 24 h timepoint).

### Analysis of gene expression at HIV transmission sites

To determine expression levels of SLC35A2 and the HIV coreceptors at different anatomical compartments, we analyzed a previously published bulk RNA-seq dataset that assessed CD4^+^ T cell populations isolated from the blood, endocervix, ectocervix, and vaginal tract of four individuals^26^. Specifically, this study gated on Live, CD45+CD3+CD4+CCR7-cells to isolate CD4^+^ T cells and further separated cells into CD103-CD69-, CD103-CD69+, and CD103+CD69+ populations before performing RNA-seq, as CD103 and CD69 are markers of resident memory T cells. We accessed raw counts from this dataset via GEO (GSE163260) and calculated counts per million (CPM) using the cpm function in the Bioconductor edgeR package^42^. CPMs were averaged across CD4^+^ T cell populations within a tissue site for each donor to determine gene expression levels in bulk CD4^+^ T cells at each site. SLC35A2 expression levels did not vary between T cell subsets within anatomical sites.

## Data availability

Illumina sequencing reads from HIV-CRISPR screens can be accessed via the SRA accession number TBD (*SRA submission in progress*). Any additional information required to reanalyze the data reported in this paper is available upon request.

## ACKNOWLEDGMENTS

We thank the Fred Hutch Shared Resources Genomics and Bioinformatics Cores, particularly Alyssa Dawson and Elizabeth Jensen for performing Illumina and Sanger sequencing, respectively; the Fred Hutch Center for Data Visualization, especially Michael Zager, Nathan Thorpe, and Sam Minot, for their support with MAGeCK analysis; Michael Emerman, Molly OhAinle, Caitlin Stoddard, and Alex Willcox for helpful discussions and technical assistance. Figures 1A, 3A, and 3D were generated with BioRender. This work was supported by the following NIH grants: NICHD R01 HD103571 to J.O. and NIGMS T32 GM007270 and NIAID F31 AI165168 to H.L.I.

## AUTHOR CONTRIBUTIONS

J.O. conceived the project and J.O., H.L.I., and D.H. designed the methodology of the study. H.L.I., D.H., and N.E.B. performed experiments. H.L.I. and D.H. conducted the data analysis, with the supervision of J.O. H.L.I. generated data visualizations. J.O. and H.L.I. wrote the paper with input from D.H. and N.E.B.

## MATERIALS & CORRESPONDENCE

Further information and requests for data, resources, and research materials should be directed to Julie Overbaugh.

